# Evaluation of behavioral and neurochemical changes induced by carbofuran in zebrafish (*Danio rerio*)

**DOI:** 10.1101/2024.01.30.578041

**Authors:** Giovana R Oliveira, Matheus Gallas-Lopes, Rafael Chitolina, Leonardo M Bastos, Stefani M Portela, Thailana Stahlhofer-Buss, Darlan Gusso, Rosane Gomez, Angela TS Wyse, Ana P Herrmann, Angelo Piato

## Abstract

Carbofuran (CF) is a carbamate class pesticide, widely used in agriculture for pest control in crops. On the other hand, CF is also detected as a contaminant in food and water sources. This pesticide has high toxicity in non-target organisms, and its presence in the environment poses a threat to the ecosystem. Research has revealed that this pesticide acts as an inhibitor of acetylcholinesterase (AChE), inducing an accumulation of acetylcholine in the brain. Consequently, this leads to the emergence of a cholinergic syndrome and various detrimental effects on human health. Nonetheless, our understanding of CF impact on the central nervous system remains elusive. Therefore, this study aimed to investigate the effects of CF on behavioral and neurochemical parameters in adult zebrafish. The animals were exposed for 96 hours to different concentrations of CF (5, 50, and 500 µg/L) and subjected to behavioral assessments in the novel tank test (NTT) and social preference test (SPT). Subsequently, they were euthanized, and their brains were used to evaluate neurochemical markers associated with oxidative stress and AChE levels. In the NTT and SPT, CF did not alter the evaluated behavioral parameters. Furthermore, CF did not affect the levels of AChE, non-protein sulfhydryl groups, and thiobarbituric acid reactive species in the zebrafish brain. However, further research is needed regarding the environmental exposure of this compound to non-target organisms.

## Introduction

Environmental contaminants such as pesticides, drugs, and heavy metals are a global concern and pose a risk to the ecosystem and public health (Caioni et al. 2021). Pesticides are used in agricultural management, pest, and disease control to increase productivity and profit. However, the widespread use of these compounds has raised numerous questions due to their potential to affect non-target organisms in the ecosystem, including humans (Dutra and Ferreira 2017; Neves et al. 2020). Contamination of food with these compounds can occur at different stages of production, such as during cultivation, processing, packaging, transportation, and storage. These molecules also have various ways of contaminating non-target organisms, such as through contact via water, food, or air (Xiong et al. 2023), which makes humans an easy target for contamination, as we are exposed throughout these processes. Contamination becomes more pronounced when considering the occupational exposure of rural workers who are directly and daily exposed to these chemicals (Ong-Artborirak et al. 2022). The increase in pesticide production and its widespread use over the last decade has resulted in a significant public health problem due to exposure to these agrochemicals, as only a minority of these compounds remain retained in the soil or are degraded; a considerable portion ends up contaminating water and food (Brovini et al. 2023). The extent of this contamination varies according to the physical and chemical properties of each compound (such as half-life, stability, and combination with other compounds that act as carriers) and soil characteristics (such as microbiota, pH, porosity, water concentration, leaching process, compound dissipation, and soil temperature) (Vryzas 2018; Mishra et al. 2020; Hu et al. 2022).

Carbofuran (CF) is a widely used carbamate class pesticide that has been detected in rivers worldwide. In Brazil, the detection of CF ranged from 1.40 µg/L to 148 µg/L (Grützmacher et al. 2008; Brovini et al. 2023) and, in China, from 1.54 µg/L to 204 µg/L (Zhang et al. 2016). The highest concentration found in Kenya was 495 µg/L (Otieno et al. 2010). In food, CF has been found above the maximum limit in cucumbers (0.146 µg/kg) (Song et al. 2018), nectar, and rapeseed pollen (35.78 µg/kg) (Wen et al. 2021), and in peanuts in Cameroon (0.0966 µg/kg) (Galani et al. 2020). CF presents high environmental persistence due to its physicochemical characteristics. It has a half-life of 36 days in water and 75 days in soil, making it resistant to degradation. Additionally, it is highly soluble in water (351 mg/dL), has high mobility in various types of soil, and has a low adsorption coefficient (Koc=30), enhancing its contamination potential in aquatic environments. It also remains stable in acidic or neutral pH soils, further contributing to its persistence in the environment (Mishra et al. 2020; Tang et al. 2021; Brovini et al. 2023). CF can be absorbed through oral, respiratory, and dermal routes in non-target organisms, causing a cholinergic syndrome, leading to symptoms such as nausea, vomiting, weakness, pain, and, in severe cases, coma and death (Mishra et al. 2020).

In preclinical studies, exposure to CF induced various behavioral and biochemical alterations. In zebrafish, exposure to CF induced anxiogenic-like effects and an increase in the activity of tyrosine hydroxylase (Liu et al. 2020). Furthermore, exposure to CF in rice fields and laboratory conditions induced oxidative damage in carp (Cyprinus carpio), leading to alterations in superoxide dismutase, catalase, glutathione S-transferase levels, and thiobarbituric acid reactive species (TBARS) levels (Clasen et al. 2014). Additionally, the impacts of CF exposure on sea bass (Dicentrarchus labrax) included decreased swimming speed and inhibition of acetylcholinesterase (AChE) (Hernández-Moreno et al. 2011). Studies on CF exposure in humans have shown an association with the development of neurodegenerative diseases (Kamel et al. 2007; Tang et al. 2021), alterations in sperm quality, and testosterone levels (Abell et al. 2000). The literature has explored the relationship between exposure to carbamates and mental disorders such as depression and anxiety (Freire and Koifman 2013).

Taking into account the problem of the widespread use of CF and its toxic potential, this study aims to evaluate the effects of CF exposure on behavioral and neurochemical parameters in adult zebrafish.

## Materials and Methods

### Animals

Experiments were performed using 160 adult (4–6 months old) short-fin wild-type zebrafish (Danio rerio, Hamilton, 1822) (50:50 male:female ratio) obtained from a local commercial supplier. Fish were acclimated to the laboratory conditions for at least 14 days before any procedure. The animals were housed in a maximum density of two fish per liter of water in 16-L unenriched tanks (40×20×24 cm) and under a 14–10-h light/dark cycle (lights on at 7 am and off at 9 pm). The tank water met controlled conditions required for zebrafish (26 °C ± 2 °C; pH 7.0 ± 0.3; dissolved oxygen at 7.0 ± 0.4 mg/L; total ammonia <0.01 mg/L; total hardness at 5.8 mg/L; alkalinity of 22 mg/L CaCO_3_; conductivity of 1500–1600 μS/cm) and was consistently aerated and filtered through mechanical, biological, and chemical filtration systems. Fish were fed twice a day with commercial flake food (Poytara®, Brazil) and brine shrimp (Artemia salina). After the behavioral tests, the animals were euthanized by hypothermic shock followed by decapitation, according to the AVMA Guidelines for the Euthanasia of Animals (Leary et al. 2020). Briefly, animals were exposed to chilled water at a temperature between 2 and 4 °C for at least 5 min after the loss of posture and cessation of opercular movements, followed by decapitation as a second step to ensure death. Following euthanasia, the sex of each animal was confirmed by dissection of gonadal tissue. For all experiments, no sex effects were observed, so data were pooled together. All procedures were approved by the animal welfare and ethical review committee at the Universidade Federal do Rio Grande do Sul (UFRGS) (approval #42114/2022).

### Chemicals

Carbofuran was obtained from Sigma-Aldrich™ (CAS 1563-66-2) (St. Louis, MO, USA). Reagents used for biochemical assays were obtained from Sigma Aldrich (St. Louis, MO, USA): bovine serum albumin (CAS Number: 9048-46-8), 5,5′-dithiobis (2-nitrobenzoic acid) (DTNB) (CAS Number: 69-78-3), thiobarbituric acid (TBA) (CAS Number: 504-17-6), and trichloroacetic acid (TCA) (CAS Number: 76-03-9). Absolute ethanol (CAS Number: 64-17-5) was obtained from Merck KGaA (Darmstadt, Germany).

### Experimental design

The animals were divided into the following experimental groups: control (CTRL, n=40) and CF (5, 50, and 500 µg/L, n=40). The concentrations were determined using environmental concentrations that had been previously reported. (Otieno et al. 2010; Zhang et al. 2016; Liu et al. 2020; Brovini et al. 2023). Allocation to experimental groups followed block randomization procedures to counterbalance sex and two different home tanks. The animals of the different experimental groups were allocated in the same room under the same conditions of temperature, luminosity, and vertical level. The experiments were carried out in two batches, with half the sample size in each batch.

During the drug exposure period, the fish were kept in 4-L static tanks (18×18×18 cm), with two tanks for each concentration to minimize potential tank effects, and remained there for 96 consecutive hours following the OECD test guidelines for the acute toxicity of substances in fish (OECD 2019). Tanks were coded to keep caregivers blinded to the treatment group and barriers were set to visually isolate the animals from different groups. CF was dissolved in ultrapure water to obtain a 40 mg/mL stock solution, which was further used to prepare the diluted solutions during the exposure. The final test concentrations were obtained by adding different amounts of the stock solution to the home tank water.

After 96 h of exposure, a set of animals was individually submitted to the novel tank test (NTT) (n=24), and a different set of animals to the social preference test (SPT) (n=16). The order of outcome assessment followed block randomization procedures to counterbalance for treatment groups. Animal behavior was video recorded (Logitech® C920 HD webcam) and analyzed by blinded researchers using ANY-Maze™ tracking software (Stoelting Co., Wood Dale, IL, USA). All tests were performed between 08:00 and 12:00 a.m. (during the light phase). Immediately after the tests, the animals were euthanized, and the brains were dissected and homogenized for the neurochemical assays. The brains of the animals submitted to NTT were prepared for analyses of oxidative stress, with a pool of 4 brains per sample. After the SPT, the animals were euthanized and the brains of these animals were collected and used to analyze the AChE activity. The neurochemical parameter analyses were as follows: non-protein sulfhydryl groups (NPSH) (n=6), TBARS (n=6), and acetylcholinesterase (AChE) activity (n=16). The experimental design is illustrated in figure 1.

**Figure 1.**
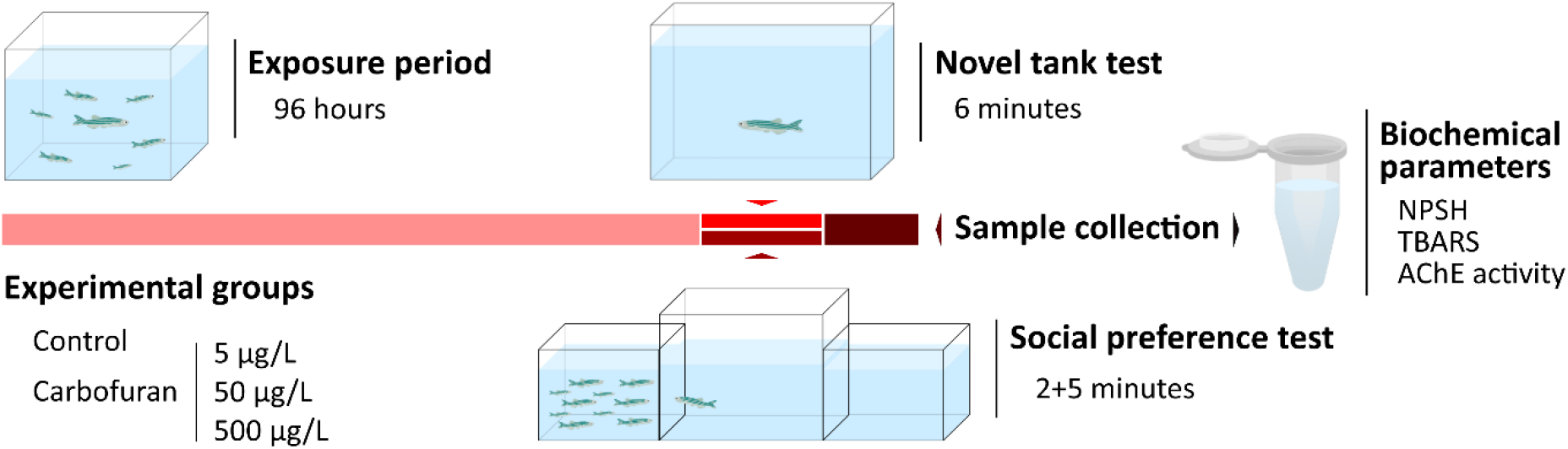
Overview of the experimental design evaluating the effects of acute carbofuran exposure to zebrafish. NPSH = Non-protein sulfhydryl groups; TBARS = Thiobarbituric acid reactive species; AChE = Acetylcholinesterase.

We report how we determined our sample size, all manipulations, and all measures in the study. Raw data were deposited in the Open Science Framework and are openly available at osf.io/2dez8/ (Rolim et al. 2023).

### Novel Tank Test (NTT)

The NTT was conducted as described previously (Marcon et al. 2018; Mocelin et al. 2019; Bertelli et al. 2021). Animals were individually placed in the NTT apparatus (24×8×20 cm and filled with water at the optimal conditions at a level of 15 cm) and recorded for 6 min. The water in the tanks was changed between animals to avoid interference from drug traces or alarm substances released by previously tested fish. The software ANY-maze™ was used to virtually divide the tank into three equal horizontal zones and track the movement of the animals. The parameters quantified were total distance traveled, mean speed, line crossings (transitions between the zones of the tank), time spent and the number of entries in the upper zone of the tank, and time spent and the number of entries in the bottom zone of the tank.

### Social Preference Test (SPT)

The SPT was conducted as described previously (Benvenutti et al. 2021; Giongo et al. 2023). In the SPT, fish were placed individually in a central tank (30×10×15cm) flanked by two identical tanks (15×10×13cm) and filmed from a frontal view for 7 min. One of the two tanks positioned beside the central tank (test tank) contained only water (neutral stimulus), and the other contained 10 zebrafish (social stimulus). All tanks were filled with water at a level of 10 cm and in the same conditions. The side of the social stimulus tank was counterbalanced to avoid any eventual bias. The water in the tanks was changed between animals to avoid interference from drug traces or alarm substances released by previously tested fish. The analyses were carried out with ANY-Maze™ software with the test tank virtually divided into three equal zones (interaction, middle, and neutral). The interaction zone was next to the tank that contained the social stimulus, while the neutral zone was considered to be next to the neutral stimulus. Animals were placed in the middle zone and had 2 min to habituate to the tank test. After this, the behavior was analyzed for 5 min. The parameters quantified were time and entries in the interaction zone, distance traveled, mean speed, and number of crossings (transitions between the zones of the tank).

### Biochemical Parameters

Following the behavioral tests, brain samples were collected according to previously published protocols (Sachett et al. 2020c). For each independent sample, four brains were collected right after the euthanasia, pooled, and homogenized in 600 μL of phosphate-buffered saline (PBS). The mixture was centrifuged at 3000 g at 4 °C in a cooling centrifuge and the supernatant was collected, which was kept in microtubes frozen at -80°C until the assays were performed (average of 2 months of storage before the analyses). The protein content was quantified according to established protocols (Sachett et al. 2020a).

### Non-protein Sulfhydryl Groups (NPSH)

The quantity of NPSH in the samples was determined following protocols optimized for zebrafish brain tissue (Sachett et al. 2021). Brain tissue preparation (50 μg of proteins) and trichloroacetic acid (TCA, 6%) were mixed. Then, it was centrifugated (10,000 g, 10 min at 4° C), and the supernatants were added to potassium phosphate buffer (TFK, 1 M). After that, the mixture was added to 5,5′-dithiobis-(2-nitrobenzoic acid) (DTNB, 10 mM) and the absorbance of 5-thio-2-nitrobenzoic acid (TNB) formed was analyzed at 412 nm after 1 h.

### Thiobarbituric Acid Reactive Species (TBARS)

Lipid peroxidation was evaluated by analyses of the production of TBARS (Sachett et al. 2020b). Samples (50 μg of proteins) were mixed with thiobarbituric acid (TBA, 0.5%) and trichloroacetic acid (TCA, 20%). The mixture was heated at 100 °C for 30 min. The absorbance was determined at 532 nm in a microplate reader. Malondialdehyde (MDA, 2 mM) was the standard.

### Acetylcholinesterase (AChE) Activity

The AChE activity was carried out using the Elmann method (Ellman et al. 1961). This technique is based on determining the thiocholine production rate. The substrate acetylthiocholine iodide is hydrolyzed by the enzyme releasing thiocholine and acetate. Thiocholine reacts with the 5,5’-dithiobis-2-nitrobenzoate ion (DTNB) to produce the yellow 5-thio-2-nitrobenzoate anion whose absorbance is detected at 412 nm in the spectrophotometer. Samples (15 μL) are pipetted into a microplate along with distilled water (220 μL) and DTNB (33 μL). Subsequently, they are incubated in a water bath at 25ºC for 3 minutes, 32 μL of 8 mM acetylthiocholine is added and absorbance is read in a plate reader at 412 nm, every 30 seconds for 2 minutes.

### Statistical Analysis

Sample size was estimated to detect an effect size of 0.5 and 0.4, respectively to NTT and SPT, with a power of 0.9 and an alpha of 0.05 using G*Power 3.1.9.7 for Windows. The total sample size was 160, equivalent to n=24 animals per experimental group in NTT and n=16 animals per experimental group in SPT. The total distance traveled was defined as the primary outcome in the NTT and the time in the interaction zone was defined as the primary outcome in SPT. The normality and homogeneity of variances were confirmed for all datasets using D’Agostino-Pearson and Levene tests. The data were analyzed using one-way ANOVA followed by Tukey’s post hoc test when appropriate. Data were analyzed and the graphs were plotted using GraphPad Prism version 8.0.1 for Windows. No outliers were excluded from the behavioral analysis data. Two data points were excluded from the control group, one from the carbofuran 5 μg/L group, two from the carbofuran 50 μg/L group, and one from the carbofuran 500 μg/L group in the AChE analysis due to discrepant results, likely arising from analytical errors. Data are expressed as mean ± standard deviation of the mean (SD). The level of significance was set at p<0.05.

## Results

### Behavioral Parameters

Figure 2 summarizes the acute effects of CF on adult zebrafish behavior in the NTT. At the tested concentrations, CF exposure did not induce alterations in the distance traveled (F_3,92_=0.005784, p=0.9994; figure 2A), line crossings (F_3,92_=0.4002, p=0.7531; figure 2B), time in the upper zone (F_3,92_=0.4235, p=0.7366; figure 2C), and entries in the upper zone (F_3,92_=0.5735, p=0.6339; figure 2D). Exposure to CF also did not elicit changes in the mean swimming speed (Suppl. figure 1A), time in the bottom (Suppl. figure 1B), and entries to the bottom zone (Suppl. figure 1C).

**Figure 2.**
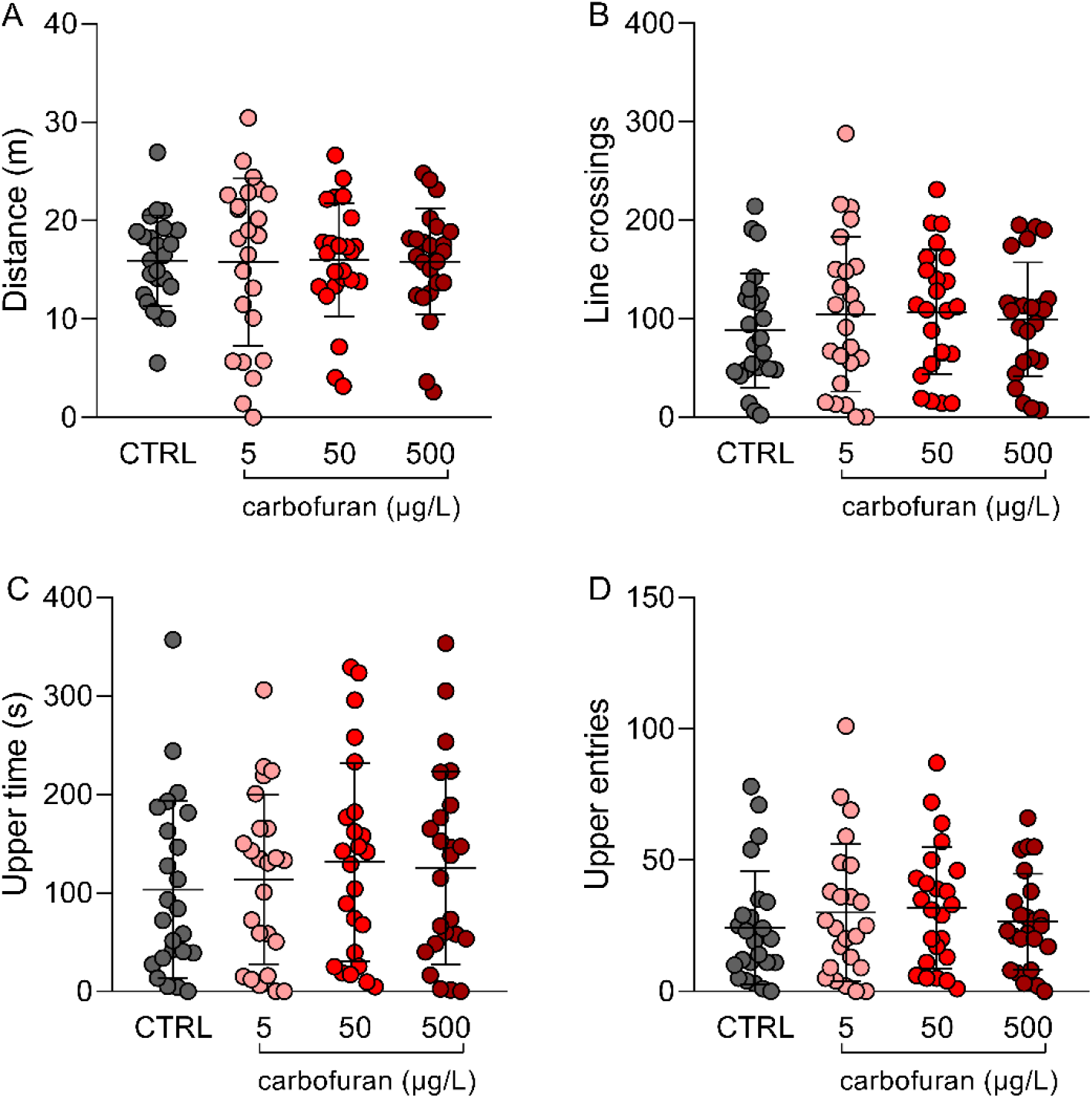
Effects of carbofuran (5, 50, and 500 μg/L) on behavioral parameters in the Novel Tank Test. (A) Distance, (B) Line crossings, (C) Upper time, (D) Upper entries. Data are expressed as mean ± S.D. One-way ANOVA. n=24. CTRL = Control.

The acute effects of CF on adult zebrafish in the SPT are shown in figure 3. CF, in the tested concentrations, did not alter the time spent in the interaction zone (F_3,60_=0.8752, p=0.4591; figure 3A), entries to the interaction zone (F_3,60_=1.315, p=0.2780; figure 3B), and distance traveled (F_3,60_=2.320, p=0.0844; figure 3C). Changes in the mean swimming speed (Suppl. figure 2A) and line crossings (Suppl. figure 2B) were also not observed. Table 1 summarizes the one-way ANOVA analysis results.

**Table 1.**
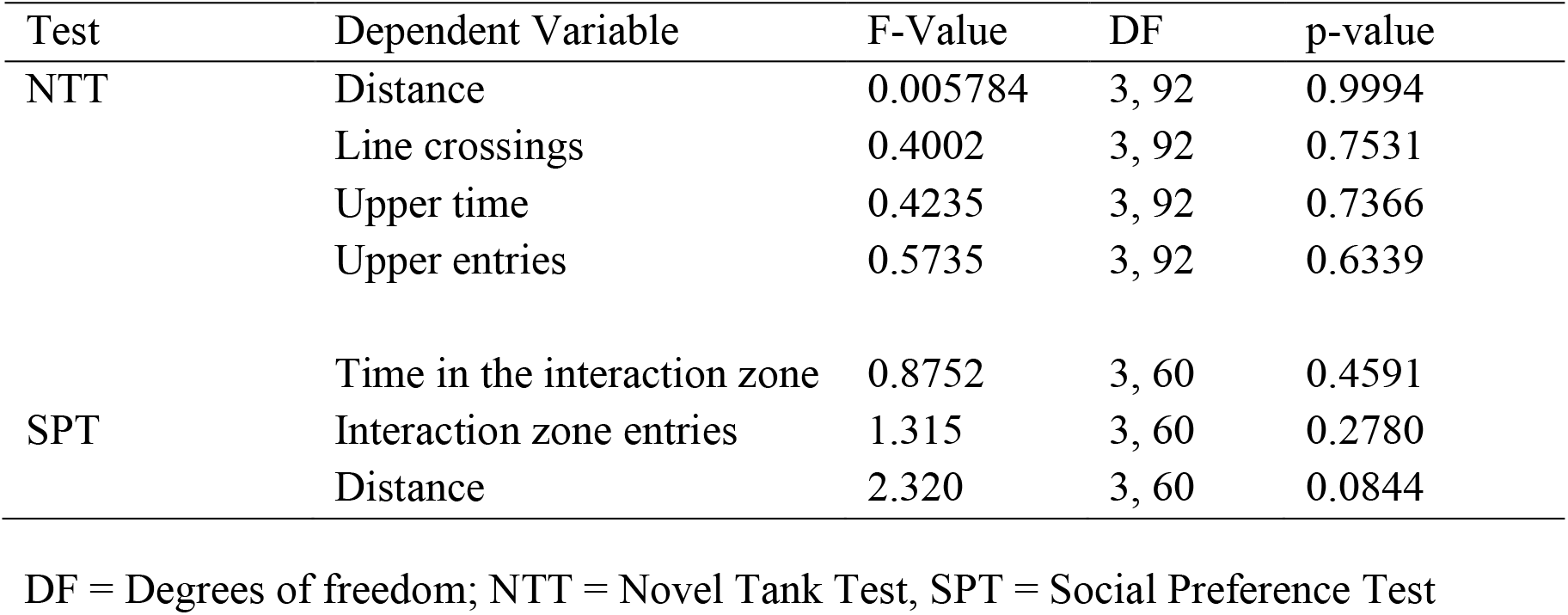
Summary of the one-way ANOVAs for the NTT and SPT.

**Figure 3.**
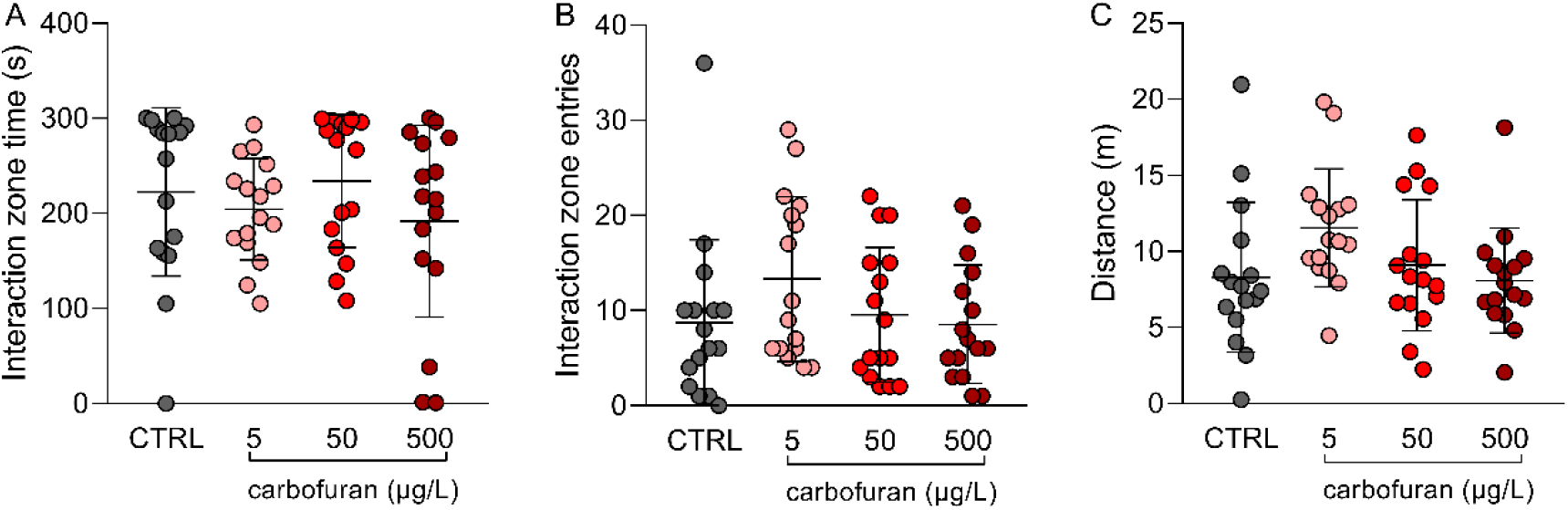
Effects of carbofuran (5, 50, and 500 μg/L) on behavioral parameters in the Social Preference Test. (A) Interaction time, (B) Interaction entries, and (C) Distance. Data are expressed as mean ± S.D. One-way ANOVA. n=16. CTRL = Control.

### Biochemical Parameters

The effects of acute CF exposure on oxidative status parameters and AChE activity in zebrafish are presented in figure 4. The exposure did not induce alterations to the levels of NPSH (F_3,20_=0.9166, p=0.4507; figure 4A) and TBARS (F_3,20_=0.4635, p=0.7109; figure 4B), and to the activity of AChE (F_3,54_=0.5667, p=0.6393; figure 4C). Table 2 summarizes the one-way ANOVA analysis results.

**Table 2.** Summary of the one-way ANOVAs for the neurochemical analyses.

| Test | F-Value | DF | p-value |
| --- | --- | --- | --- |
| NSPH | 0.9166 | 3, 20 | 0.4507 |
| TBARS | 0.4635 | 3, 20 | 0.7109 |
| AChE activity | 0.5667 | 3, 54 | 0.6393 |

DF = Degrees of freedom; NPSH = Non-protein sulfhydryl groups; TBARS = Thiobarbituric acid reactive species; AChE = Acetylcholinesterase.

**Figure 4.**
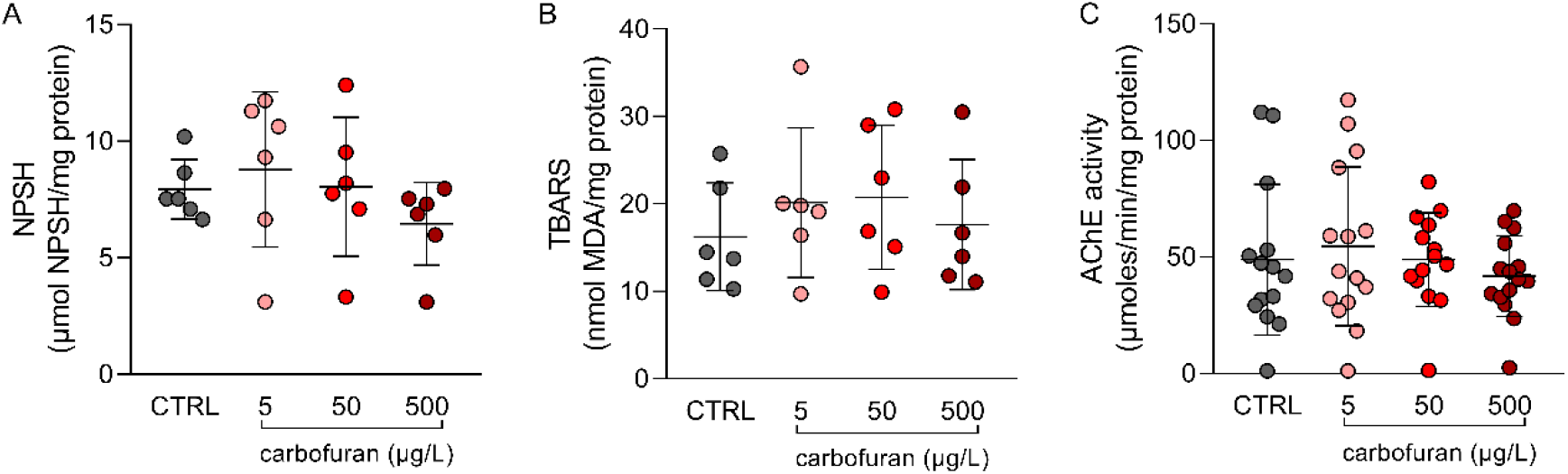
Effects of carbofuran (5, 50, and 500 μg/L) on neurochemical parameters. (A) NPSH (Non-protein sulfhydryl groups), (B) TBARS (Thiobarbituric acid reactive species), (C) AChE (Acetylcholinesterase) activity. Data are expressed as mean ± S.D. One-way ANOVA. n=6 for NPSH and TBARS, n=14-15 for AChE (n=14 CTRL; n=15 CF5; n=14 CF50; n=15 CF500).CTRL = Control.

## Discussion

Here, carbofuran (at concentrations of 5, 50, and 500 μg/L) showed no impact on zebrafish behavior during both the novel tank and social preference tests. Moreover, there were no alterations observed in the analyzed neurochemical parameters, such as NPSH, TBARS, and AChE activity.

The novel tank test (NTT) is a well-established tool for assessing the effects of anxiety-like behavior modulating interventions in zebrafish (Kysil et al. 2017; Johnson et al. 2023). It capitalizes on the natural behavior exhibited when these fish encounter a novel environment. Zebrafish typically display an innate preference for the bottom zone, a behavior that gradually shifts after habituation to the new setting, leading them to explore other zones of the tank (Cachat et al. 2010). Anxiogenic interventions tend to exacerbate the zebrafish’s initial preference for the bottom region, amplifying anxiety-like behavior (Kysil et al. 2017). Conversely, anxiolytic drugs or interventions that disrupt their innate defensive behavior prompt fish to spend more time in the upper zone (Gebauer et al. 2011; Kysil et al. 2017). Additionally, total distance traveled serves as a key metric for overall locomotor activity, offering valuable insights into the hyperstimulating or sedating effects of various interventions (Cachat et al. 2010; Kysil et al. 2017). The locomotor behavior of this species is a complex behavior controlled by the activity of various brainstem neurons along with neuromodulatory projections whose activities depend on the integrity of these cells of the animal (Drapeau et al. 2002; Dreosti et al. 2015; Sachett et al. 2022). The exploratory, locomotor, and anxiety-related parameters studied in the novel tank test are essential for reproduction, predator avoidance, and food-seeking behaviors (Zhang et al. 2016). Notably, a study assessing the impact of exposure to the herbicide 2,4-D on zebrafish has observed alterations in both the time spent in the upper zone and the distance traveled, disrupting their natural behavior (Thiel et al. 2020). Furthermore, a recent meta-analysis showed that fungicides decrease the locomotion in both zebrafish larvae and adults (Reis et al. 2023). In our study, there is no evidence that CF alters any locomotor parameter.

The social preference test (SPT) stands as a valuable test for investigating the effects of various interventions on zebrafish, given their innate schooling behavior and intricate social dynamics, including hierarchical and breeding relationships (Dreosti et al. 2015). Being a species with complex social interactions, zebrafish engage in social behaviors as a means of escaping predation and to facilitate reproduction (Dreosti et al. 2015). Given the critical role of social preference in this species, alterations in this behavior are highly relevant, making the SPT an ideal assay for studying interventions that may disrupt the natural behavior of zebrafish. Despite findings from previous studies indicating alterations in social behavior due to exposure to different pesticides (Yan et al. 2023; Liu et al. 2023), our study did not observe any changes following exposure to CF in the SPT. Since social behavior is genetically preserved and has ontogenic nature (Buske and Gerlai 2011; Scerbina et al. 2012), it is less vulnerable to low-impact modulations, such as the concentrations used in this study, where the highest concentration corresponds to 6% of the LC_50_ of similar species, such as Carassius auratus, the goldfish (Dreosti et al. 2015; Valadas et al. 2023).

Reactive oxygen species are constantly produced during normal physiological events and are important in the brain signaling processes. High concentrations of oxidants can damage cell components, including lipids, and such damage leads to physiological damage with impairment of normal cell functions. Neuronal membranes are rich in polyunsaturated fatty acids, which make the brain more vulnerable to lipid oxidation and to the damage caused by lipid peroxidation (Fedoce et al. 2018; Halliwell 2022). Within this study, there were no changes in the presented markers of oxidative stress and cellular damage. However, other oxidative stress markers of interest have been elucidated in carbofuran exposure, such as SOD and CAT enzymes (Fedoce et al. 2018; Liu et al. 2020). Another crucial aspect to consider when discussing the results found here with those already reported in the literature is the animal experimentation protocol. This study followed OECD 203 (OECD 2019), a guideline that regulates and standardizes toxicological exposure studies on zebrafish. This guideline advocates exposing the animals for 96 hours before commencing behavioral tests, avoiding the use of solvents in compound dilution, and avoiding any disturbance that may alter the animal’s behavior. Several studies presented here did not follow these guidelines, such as the study conducted by Liu and collaborators (Liu et al. 2020), which exposed the animals for 48 hours and handled the animals to replace the aquarium water, which is a stressful factor for the animals and could be associated with some of the behavioral changes reported, as well as the studies by other research groups that also did not follow this standardized protocol (Hernández-Moreno et al. 2011; Clasen et al. 2014; Cui et al. 2019; Mendes et al. 2021).

Carbamates, including CF, act as reversible inhibitors of AChE, causing carbamylation of this enzyme. This carbamylation prevents the degradation of acetylcholine, allowing for its prolonged action in the synaptic cleft. Consequently, it leads to hyperexcitability of both nicotinic and muscarinic cholinergic receptors, resulting in cholinergic syndrome (Gupta 1994; Mishra et al. 2020). CF has consistently been shown to inhibit AChE in different fish species (Dembélé et al. 2000; Hernández-Moreno et al. 2011). Here, on the other hand, no alteration in the activity of AChE was measured after exposure to CF. This could be attributed to the enzymatic reactivation of AChE. The zebrafish’s brain regeneration after injury is complex and has several repair mechanisms, as previously reported, which could be associated with the adaptation to the damage caused by carbofuran after AChE reactivation and normalization of enzymatic activity (Ghosh and Hui 2016).

An important aspect to be discussed is the association of carbofuran with other compounds. Carbofuran has been detected in various samples together with chlorpyrifos, 2,4-D, glyphosate, and other pesticides of great toxicological importance (Buch et al. 2013; Brovini et al. 2023), which could suggest an association and interaction between these compounds. A study conducted in 2016 reports the interaction between carbofuran and carbon nanotubes, where the association of these two pollutants increases histological damage in Nile tilapia (Oreochromis niloticus) exposed to both compounds by 25% compared to animals exposed only to carbofuran (Campos-Garcia et al. 2016). Carbofuran exposure in association with carbon nanotubes also significantly affected the metabolic route of shrimp (Palaemon pandaliformis) (Alves et al. 2022). Carbofuran combined with cadmium-induced acute synergistic effects, increased oxidative stress, endocrine effects, and increased cellular apoptosis in adult and larval zebrafish (Wang et al. 2022). Regarding the formulation, the one used in this study was the isolated compound without the use of a commercial formula. In the environment, the pesticide is used in a commercial formulation, where the active ingredient is mixed with so-called inert compounds that comprise more than half of the product’s volume. A study conducted on aquatic species demonstrates differences in toxicity between carbofuran and Furadan® (its commercial formulation) as well as the potential interaction of carbofuran with compounds like diuron that can cause different toxic responses than predicted for individual compounds (Mansano et al. 2016). Regarding the mixture of Furadan® and RoundUp® (Glyphosate, commonly found in association with carbofuran), an increase in glycoproteins-P was observed, and even with concentrations below the permitted levels, the combination of the two compounds presented cytotoxicity in the ZFL cell line (a well-established zebrafish hepatocyte cell line) (Goulart et al. 2015), suggesting the toxicity of surfactants present in the formulation, which, when combined with pesticide compounds, enhance harmful effects.

## Conclusion

Although the results presented in this study do not show behavioral and neurochemical changes, it is still necessary to study the effects of carbofuran under conditions closer to the real environment where the pesticide is present and causing harm to non-target organisms. To further contribute to the fields of toxicology and the environment, it is essential to study a broader range of concentrations of this compound and the alterations it induces in different outcomes. Additionally, studying longer exposure times to environmental concentrations and their transgenerational and long-term effects at different stages of zebrafish is crucial.

## Author Contributions

Conceptualization: G.R.O., M.G.-L., and A.P. Data curation: G.R.O. and M.G.-L. Formal analysis: G.R.O., M.G.-L., and A.P. Funding acquisition: A.P. Investigation: G.R.O., M.G.-L., R.C., L.M.B., S.M.P., T.S.-B., and D.G. Methodology: G.R.O., M.G.-L., R.C., D.G., A.T.W., A.P.H., and A.P. Project administration: G.R.O., M.G.-L., and A.P. Resources: A.T.W. and A.P. Supervision: A.P. Validation: G.R.O., M.G.-L., and A.P. Visualization: G.R.O., M.G.-L., and A.P. Writing - original draft: G.R.O. and M.G.-L. Writing - review & editing: R.C., L.M.B., S.M.P., T.S.-B., D.G., R.G., A.T.W., A.P.H., and A.P.

## Interest Statement

The authors declare no conflicts of interest.

## Data Availability

All data are available in Open Science Framework (osf.io/2dez8/).

## Acknowledgments

The authors thank the Conselho Nacional de Desenvolvimento Científico e Tecnológico (CNPq, proc. 303343/2020-6), Coordenação de Aperfeiçoamento de Pessoal de Nível Superior - Brasil (CAPES), Instituto Nacional Saúde Cerebral (INSC, No. 406020/2022-1)/CNPq, and Pró-Reitoria de Pesquisa (PROPESQ) at Universidade Federal do Rio Grande do Sul (UFRGS) for funding and support.

## Notes

### Competing Interest Statement

The authors have declared no competing interest.

