## Supplementary Material for "Evaluation of behavioral and neurochemical changes induced by carbofuran in zebrafish (*Danio rerio*)"

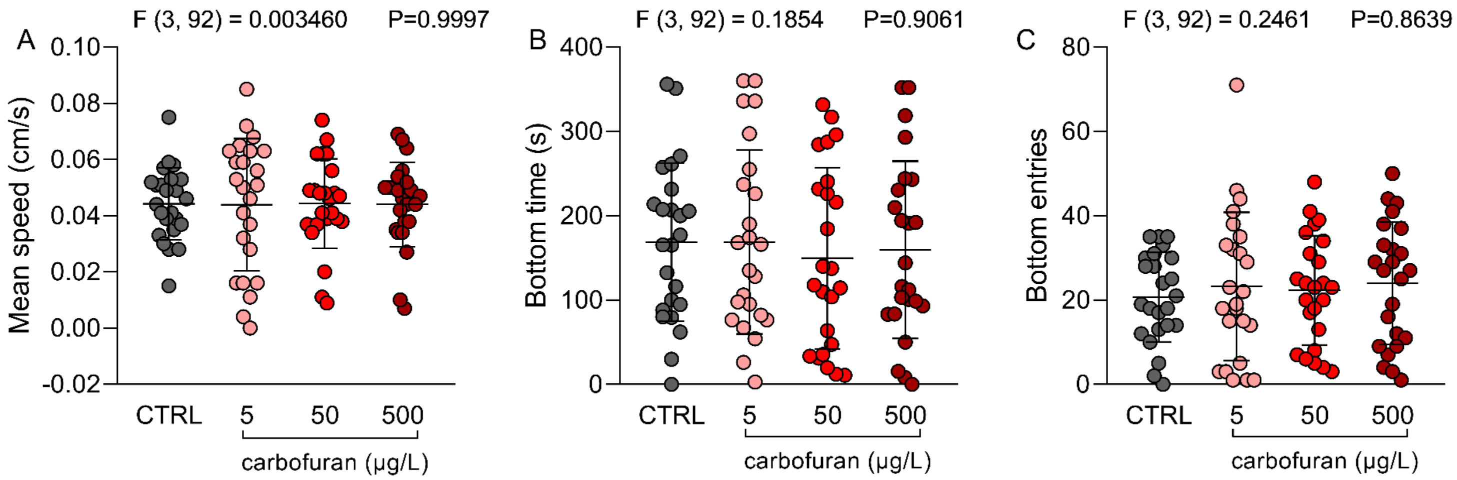


**Suppl. figure 1.** Effects of carbofuran (5, 50, and 500 µg/L) on behavioral parameters in the Novel Tank Test. (A) Mean speed, (B) Bottom zone time and (C) Bottom zone entries. Data are expressed as mean ± S.D. One-way ANOVA. n=24. CTRL = control

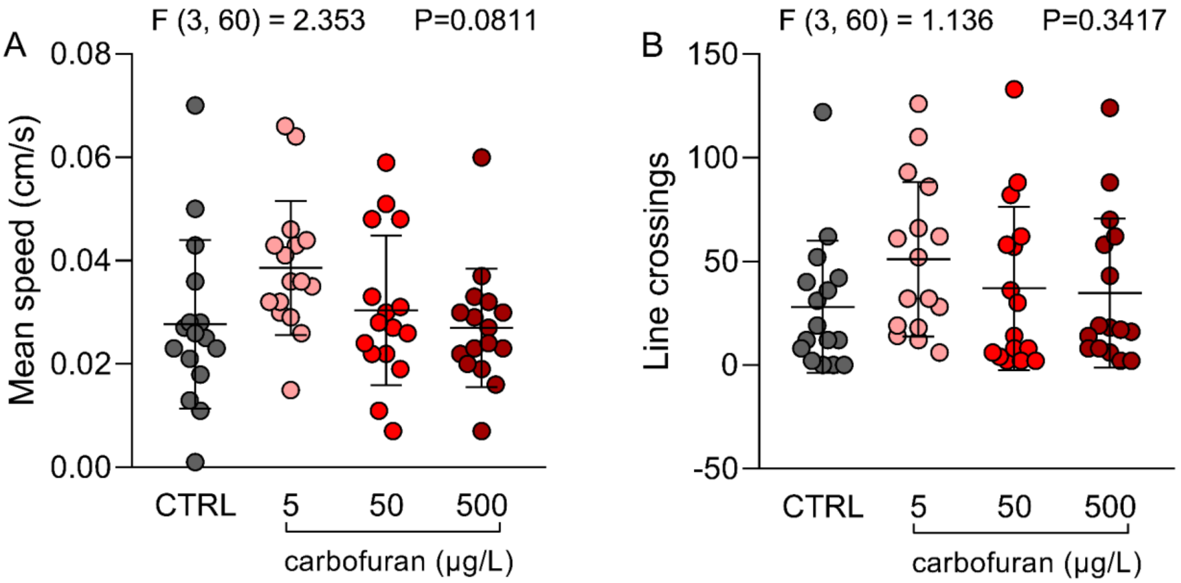


**Suppl. figure 2**. Effects of carbofuran (5, 50, and 500 µg/L) on behavioral parameters in the Social Preference Test. (A) Mean speed, (B) Line crossings. Data are expressed as mean ± S.D. One-way ANOVA. n=16. CTRL = control.
